# Evolutionary conservation of secondary structures in the lncRNAs of plants

**DOI:** 10.1101/2023.08.13.553158

**Authors:** Jose Antonio Corona-Gomez, Peter F. Stadler, Selene L. Fernandez-Valverde

**Author notes:** Present address: School of Biotechnology and Biomolecular Sciences and the RNA Institute, The University of New South Wales, 2052, Sydney, NSW, Australia. Corresponding author –.

## Abstract

LncRNAs are essential regulators of eukaryotic gene expression. They exert their gene regulatory functions by interacting with DNA, RNA, and protein. These functions are considered at least in part associated with their capacity to fold into complex three-dimensional structures. The conservation of lncRNA structure in mammalian genomes has been assessed in several studies, however, very little is known about the conservation of lncRNA structures in plants. Here, we analyze the structural conservation of lncRNAs in *Brassicaceae*, using a whole genome alignment of 16 *Brassicaceae* species. We found that 44.2% (1925 of 4354) of the intergenic lncRNAs (lincRNAs) and 75.1% (1549 of 2060) of the natural antisense transcripts (NATs) of *Arabidopsis thaliana* have conserved structural motifs in at least 2 of the 16 species. Also, 3612 lncRNAs have conserved structural motifs in multiple species; 2264 of which are tissue-specific, and 841 can be associated with a function by a co-expression network in *A. thaliana*. Indeed, we find evidence for the conservation of structural motifs in several lncRNAs with known functions, including, *lncCOBRA1, FLORE, IPS1, ELENA1* and *COOLAIR.* The latter was shown previously to have a conserved structure. Overall, we have identified numerous lncRNAs with conserved structures in *Brassicaceae* that warrant further experimental exploration *in vivo* to understand whether these lncRNAs and their conserved structures are of biological significance.

## Introduction

LncRNAs are regulatory RNAs over 200 nt (Derrien et al. 2012) that have little or no coding potential. Most lncRNAs are transcribed by RNA Pol II and mature similarly to mRNAs (Mercer and Mattick 2013). In plants, two additional RNA polymerases, Pol IV and Pol V, can transcribe lncRNAs (Chekanova 2015; Wierzbicki et al. 2008; Zhang and Chen 2013). Despite their low conservation, many lncRNAs perform essential gene regulatory functions (Quinn and Chang 2016). In plants, they have been identified as participants in the modulation of chromatin topology, interaction with miRNAs (miRNA sponges), splicing modulation, small RNA precursors and scaffolds for the formation of protein complexes (Wang et al. 2014; Kim et al. 2017; Zhang et al. 2014; Franco-Zorrilla et al. 2007; Rai et al. 2019). They participate in important biological processes such as flower (Yamaguchi and Abe 2012; Jampala et al. 2021), fruit and root development (Bazin and Bailey-Serres 2015; Ariel et al. 2020), shade-avoidance response (Wang et al. 2014; Li et al. 2021), and leaf senescence, as well as in the response to various biotic and abiotic stresses (Jha et al. 2020), such as drought, salt and cold stresses.

The conservation of genes, including lncRNAs, is indicative of a character or property shared between species despite the speciation process, suggesting it might be associated with a biological function. The conservation of a gene is usually measured by comparing its sequence with orthologous genes from different species. This amounts to counting the equal columns of an alignment of nucleotides or amino acids (Hedrick and Miller 1992). Long stretches of conservation are generally useful for identifying conserved protein-coding genes. However, lncRNAs are generally poorly conserved at the sequence level compared to coding genes (Pang et al. 2006; Marques and Ponting 2009; Ponjavic et al. 2007). This lack of sequence conservation has prompted the exploration of other types of conservation, such as i) conservation by genomic position (Mohammadin et al. 2015; Nitsche and Stadler 2017; Smith et al. 2013), ii) splicing conservation (Corona-Gomez et al. 2020; Nitsche et al. 2015; Schüler et al. 2014), iii) conservation of expression (Hezroni et al. 2015; Necsulea et al. 2014), iv) functional conservation (Ulitsky et al. 2011), v) short conserved motifs (Ross et al. 2021) and vi) conservation of RNA structural folding (Seemann et al. 2017; Smith et al. 2013). Each different type of conservation in lncRNAs has helped reveal conserved sequence or structural motifs, which might be preserved due to the interactions of lncRNAs with other molecules such as lncRNA: DNA, lncRNA: RNA and lncRNA: protein (Ross and Ulitsky 2022).

RNA structures are formed by the union of ribonucleotides in the form of loops and hairpins within the same molecule (Cruz and Westhof 2009; Li et al. 2012; Stombaugh et al. 2009). Many regulatory RNAs, such as miRNAs, depend on their structure for their processing (Bartel 2004). In the case of lncRNAs, it is known that their structure can play an important role in their function, but the information is limited to some well-studied cases of all identified lncRNAs, such as the dosage-compensating lncRNAs *rox1* and r*ox2* in *D. melanogaster* (Ilik et al. 2013; Quinn et al. 2016), the RepA repeat in the human *Xist* gene (Maenner et al. 2010), and *COOLAIR,* a regulator of flowering in *A. thaliana* (Hawkes et al. 2016). There are examples of functional RNA motifs that can be readily recognized by their structure, such as riboswitches (Pedersen et al. 2006) or RNA-protein interaction motifs (Ray et al. 2013). However, these readily identifiable motifs are seldom present in the 4094 examples of RNA families in the Rfam 14.8 database; where each family is represented by a multiple sequence alignment, a consensus secondary structure, and has some associated functional classification (Kalvari et al. 2021). Only 218 of these families are exclusive to lncRNAs.

Structural conservation is a good predictor of function in biopolymers; this is observed in proteins and nucleic acids, even though they have completely different forms of folding (Fallmann et al. 2017). In *A. thaliana,* it has been proposed that around 32,000 structured RNA regions are expressed (Li et al. 2012). With this information, it can be assumed that the structures are a common element in *A. thaliana* transcripts. However, how many of these structures are under selection is a question that has not been widely addressed. We can mention a few examples of studied lncRNAs with conserved structure and function in *A. thaliana*. The most well-known example is *COOLAIR,* a NAT involved in vernalization, which has been shown to have a complex structure conserved in *Brassicaceae* (Hawkes et al. 2016). Another example of a lncRNA with a known structure and function is *HID1*, which is involved in the photomorphogenesis process and regulates the *PIF3* gene by binding directly to its promoter region. Homologous genes to *HID1* have been found in many plant species and shown to have the same function, for example, the homologous gene from rice can be used to recover the phenotype of a *HID1* mutant in *A. thaliana (Wang et al. 2014)*.

In this article, we identified conserved RNA structures (CRSs) in lncRNAs across *Brassicaceae*. We sought to measure how many *A. thaliana* lncRNAs might be conserved. We also identified lncRNAs that are under stronger structural selection compared to other genes to help to pinpoint candidate functional lncRNAs in *A. thaliana* that might warrant further characterization.

## Results

### Number of CRSs identified

We identified a total of 28,461 CRSs in *Brassicaceae* using the locARNA-p and CMfinder methods. Interestingly, both methods predicted similar numbers of CRSs, with 20624 and 18210 predicted by locARNA-p and CMfinder, respectively, across 10,373 transcripts (Figure 3A). However, the methods differ in their approach to identifying these structures. Specifically, locARNA-p tends to identify larger CRSs while CMfinder focuses on smaller, well-conserved motifs of 200 bases (Figure S1). This results in differences in the size of the structural alignment, which in turn affects the number of paired nucleotides in the consensus structural motif. Notably, the locARNA-p method produces more paired nucleotides on average than the CMfinder method (Figure S1).

### Conservation of CRSs in lncRNAs compared to other RNAs

We analyzed a total of 6,764 lncRNAs and found that 54.4% of them (3685 lncRNAs) contain CRSs (Figure 3B). While this percentage is lower than what we observed in tRNAs (69.7%, or 480 out of 689) and mRNAs (71.1%, or 19,672 out of 27,655), it is higher than what we found in unannotated regions of the genome (26.3%) (Figure 3B). These findings suggest that CRSs are featured in all types of RNAs, including lncRNAs, though their prevalence may vary in different types of genes. Further analysis revealed that 1925 lincRNAs, 1549 NATs, 117 sense exonic lncRNAs, and 122 intronic lncRNAs contained CRSs (Figure 3C). NATs had the highest percentage of CRSs (75.1%) compared to other lncRNA subtypes, such as lincRNAs (44.2%).

### Percentage of conserved structural motifs

We discovered that 62.4% (4220/6764) of the lncRNAs in the 16 *Brassicaceae* species are conserved by genomic position. Of these, roughly 43.6% (2947/6764) contain CRSs (Figure 4). This implies that there are 2947 lncRNAs in the *Brassicaceae* family that maintain their location and exhibit a conserved secondary structure (Figure 4). This finding indicates that the location of RNA structures may be critical to their function since the majority of these structures function in CIS. However, this result is also influenced by the methodology we used to identify the CRSs in all 16 species, as we first identify the coordinates and then the structure of the 2947 lncRNAs with CRSs in all 16 species, 1409 are NATs and 1367 of the annotated lincRNAs.

### Structural conservation over sequence conservation

We observed that 29.9% of lncRNAs and 35.8% of mRNAs contain CRSs with observed covariance that is greater than expected based on phylogenetic distance, according to R-scape. However, we acknowledge that this method is stringent, and it can be challenging to differentiate between sequence and structure conservation when the phylogenetic distance is inadequate. While less than one-third of lncRNAs with CRSs displayed greater observed covariance than expected, it is important to note that the lack of covariance does not necessarily imply the absence of conserved structure, as structure conservation cannot be distinguished from sequence conservation (Rivas et al. 2020).

Previously, we identified tissue-specific lncRNAs in *A. thaliana* using *Tau* values in 24 different tissue or life stage categories (Corona-Gomez et al. 2022). Among the 4188 lncRNAs specific to each category, 2269 had CRSs. The highest number of lncRNAs with CRSs was found in the embryo (705 lncRNAs) and in the whole adult plant (402 lncRNAs), consistent with previous data showing these categories having the highest number of unique lncRNAs (Figure 5A). On average, 60.2% of lncRNAs in each of the 24 categories had CRSs. The seedling had the highest percentage of lncRNAs with CRSs at 79.8% (Figure 5A). In comparison, among mRNAs, the highest number of mRNAs with CRSs was found in the root (2029). The highest percentage of mRNAs with CRSs was found in the cotyledon and hypocotyl, with an average of 81% in the 24 categories (Figure 5A). These findings suggest that RNAs with CRSs are not uniformly distributed in all tissues. For lncRNAs, we observed a significant number of these in the embryo and a large percentage conserved in the seedling, which leads us to hypothesize that the conserved structures in lncRNAs might be important in these life stages of *A. thaliana*.

Furthermore, we identified 841 lncRNAs with CRSs present in our previously reported co-expression modules (Corona-Gomez et al. 2022). Remarkably, this represents a substantial proportion of the lncRNAs in the co-expression modules, as 67.8% (841/1241) of them contain CRSs (Figure 5B). These lncRNAs are relevant as they can provide insights into the biological processes they participate in through the guilty-by-association approach, which we summarized into functional categories in our previous work (Corona-Gomez et al. 2022). Among the functional categories, the one with the most lncRNAs with CRSs is related to chloroplast organization and photosynthesis (Figure 5B), which includes 292 lncRNAs. This category is especially relevant since it involves plant photosynthetic tissues and response to different light intensities.

### Examples of lncRNAs with conserved structural motifs

Among the lncRNAs with CRSs, We have identified four lncRNAs with known functions in the literature: *COOLAIR*, *FLORE*, *IPS1*, *lncCOBRA1,* and *ELENA1*. The first one, *COOLAIR*, is the best-characterized natural antisense transcript (NAT) of *A. thaliana* (Swiezewski et al. 2009)) and has a conserved structure in the *Brassicaceae* family that has been experimentally proven (Hawkes et al. 2016) Further, it has been demonstrated in vivo that the structure of COOLAIR can vary depending on the isoform and environmental conditions, leading to a range of structures that can modify its regulatory function (Yang et al. 2022). We identified CRSs in the second exon of *FLORE*, which is a NAT to the CDF5 coding gene.

*FLORE* promotes flowering by regulating the CDF genes involved in the circadian cycle of the photoperiod (Henriques et al. 2017). The mRNA that overlaps with *FLORE* also contains CRSs, suggesting a potential interaction between the two structures. This is consistent with previous findings that these genes are known to regulate each other, and that *FLORE* can even regulate other CDF proteins in *trans (Henriques et al. 2017)*. *IPS1* functions as a sponge for miRNA mir399 in phosphate regulation (Franco-Zorrilla et al. 2007). The region where mir399 binds to *IPS1* has a highly conserved CRS. It is known that the folding of RNAs can expose certain regions of a transcript, favoring interactions with other RNAs, including miRNAs. We thus hypothesize that the predicted structural motif of IPS1 may serve this function. The lincRNA *lncCOBRA1* contains CRSs and is known to be involved in seed germination, as well as interacting with a diverse range of proteins (Kramer et al. 2022). The identified CRS might relate to the function of *lncCOBRA1* as a protein scaffold. *ELENA1*, is involved in plant immunity and response to pathogens (Mach 2017). Although it has the potential to produce peptides, it has been demonstrated that its function is not related to peptide production due to a mutation in the start codon. Instead, *ELENA1* binds to the metering subunit 19a (MED19a) in response to pathogens (Seo et al. 2017), this binding in turn positively regulates pathogen response genes, such as PR1, by binding to its region promoter (Mach 2017; Rai et al. 2019) This interaction between the *ELENA1* RNA, protein, and chromatin may be associated with its structural motif, which was found in all 16 species of *Brassicaceae* analyzed.

Our analysis identified CRSs in four out of the six isoforms of *COOLAIR* conserved in all 16 species analyzed. The isoforms included one proximal and three distal types (Figure 6). We also found CRSs in the FLC coding gene that overlaps with *COOLAIR* (Figure 6). Our findings were consistent with the helix structures proposed by Hawkes et al. (2016) using the SHAPE technique, which allows experimental characterization of RNA structures. In the case of proximal *COOLAIR*, our prediction coincides with helix H5 and in the distal *COOLAIR* isoform, our predictions coincide with helix H3 and H12 (Figure S2). It’s important to mention the H3 helix in the distal isoform of *COOLAIR* is conserved in five species of *Brassicaceae (Hawkes et al. 2016)*, and in our results, this is present in 16 species, which tells us about the importance of this structure. Additionally, the initial 5’-m region, including helices H1, H2, and H3, had low reactivity by SHAPE, which is associated with the formation of R-Loops that are crucial for *COOLAIR* function (Xu et al. 2021; Sun et al. 2013),. Thus, the identified structure may play a role in the formation and size of R-Loops.

## Discussion

We found numerous (3685) lncRNAs with CRSs conserved in the *Brassicaceae* family, which had not been previously reported, giving a total of 75.1% of the NATs and 44.2% of the lincRNAs with CRSs. The proportion of lncRNAs with CRSs is similar to that found in vertebrates. A catalog of lncRNAs and mRNAs with CRSs was generated, which forms a basis for the search for functions of RNA structures in plants.

Our study reveals that the structural conservation of long non-coding RNAs (lncRNAs) is greater than the conservation observed by sequence or splicing (Corona-Gomez et al. 2020) but lower than conservation by genome position. To increase the number of lncRNAs analyzed, we incorporated the most recent annotation in this study (Corona-Gomez et al. 2022), resulting in a larger number of lncRNAs conserved by genome position. Our updated analysis revealed a higher proportion of lncRNAs conserved by position, and the discovery of structural conservation in most lncRNAs is novel. In comparison to previous studies, we found that plants exhibit higher conservation percentages of lncRNA CRSs than vertebrates (Seemann et al. 2017). However, it should be noted that the vertebrate study had a larger distance between the species analyzed than ours. As expected, coding genes showed the highest conservation of CRSs in both plant and vertebrate species.

Our findings (Figure 4) reveal that the conservation of lncRNAs by position and structure exceeds the 38% previously reported for non-coding genome sequences (36 bp genome regions) between the *A. thaliana* and *A. arabicum* genomes (Haudry et al. 2013). In comparison to the proportion of 3% of conserved lincRNAs previously estimated between *A. thaliana* and *A. arabicum* (Nelson et al. 2016), our results demonstrate that the percentage of conservation is almost 10 times greater by integrating structure and position conservation. The number of lncRNAs reported as homologs between *A. thaliana* and *A. arabicum* is 503 in one of the largest annotations of lincRNAs to date (Palos et al. 2022). For comparison, we found 904 lincRNAs with genome position conservation and CRSs in the 16 species analyzed. This highlights the significance of structure and position as essential factors in studying lncRNAs, underscoring their importance in advancing our knowledge of this emerging field.

Comparing our bioinformatic results against experimental structural prediction analyses in *A. thaliana*, we see that 13.11% of our predictions match the high structure peaks predicted in immature flower buds by the method dsRNA-seq and ssRNA-seq (Li et al. 2012). This method is based on the sequencing of the enzymatic digestion of RNA by means of specific ribonucleases of double or simple chain (Zheng et al. 2010); comparing the proportion of readings of both digestions, it is determined if the transcript is in a double or simple chain at the time of enzymatic digestion. The low percentage of coincidence in this comparison is in the fact that they are different methods, and the experimental methods are always limited to a single condition when coming from a single tissue or stage of development of the plant (Li et al. 2012), For this reason, it is to be expected that it does not contain all the gene annotations nor does it represent all the possible structures in them, in addition to another important factor, for example in the comparison of lncRNAs, is the depth to find them, which makes it more difficult to identify, yet we found that 282 of lncRNAs have structure presence by the dsRNA-seq and ssRNA-seq method and by our prediction.

The results of the structures must be taken considering the error rate of each method, where for the results of locARNA-p a 13% of false positives is expected (Will et al. 2012; Wilm et al. 2006) and for those of CMfinder 21% (Yao et al. 2006). Despite the error rate, lncRNAs with CRSs are abundant and are good candidates for searching functional lncRNAs given their conservation of a functional structure. Both CMfinder and locARNA-p methods agree on predicting conserved structural motifs in the same gene on average 51.5% of their predictions, but this is not distributed in the same way in the different types of RNAs: in tRNAs the percentage of genes with positive prediction in both methods is 78.4%, being the maximum and the minimum the unannotated regions with 3.3%. The factor that most influences both methods to find CRSs is gene sequence conservation, another factor that plays an important role in both methods making a simultaneous prediction is that CMfinder only considers small, well-structured motifs, which generally favors small motifs with high GC as those present in tRNAs while locARNA-p motifs with a smaller GC and larger as those characteristics of lncRNAs. To further investigate this, we plan to use more phylogenetically distant species in the future to determine whether the percentage of covariance increases with greater phylogenetic distance.

## Materials and methods

### Extraction of orthologous sequences by genomic position

We identified the orthologous positions of the genes by using our previously generated whole-genome alignment (WGA) of 16 species of the *Brassicaceae* family (Corona-Gomez et al. 2020). As input data, three types of genes were used: high-fidelity lncRNAs annotated in previous works (Corona-Gomez et al. 2022), Araport11 coding genes, and Araport11 tRNAs (Cheng et al. 2017). As a control, we also incorporated unannotated regions of the genome of 1000 bases, to extract these unannotated regions of the genome we used the annotation of Araport11 (Cheng et al. 2017) in addition to the annotation of the lncRNAs of Corona-Gomez et al. 2022 and the complement function of bedtools (Quinlan and Hall 2010), then with the random function of bedtools (Quinlan and Hall 2010) we extracted 1000 bases regions of the *A. thaliana* genome. The annotation and coordinates of the genes were taken in BED12 format since the analysis is sensitive to the introduction of introns, so these are removed, and only exons are used as a representation of mature RNAs for the analysis. The HALtools program (Hickey et al. 2013) was used to extract the orthologous positions in each species from a previously generated genomic alignment of 16 *Brassicaceae* species (Corona-Gomez et al. 2020). After this, the nucleotide sequence of each of the species was extracted using the orthologous positions and saved in fasta format (Figure 1). It is important to mention that each isoform was analyzed individually for genes with isoforms. If more than one orthologue was present in a particular species, only the copy with the highest sequence identity with the reference was analyzed.

**Figure 1.**
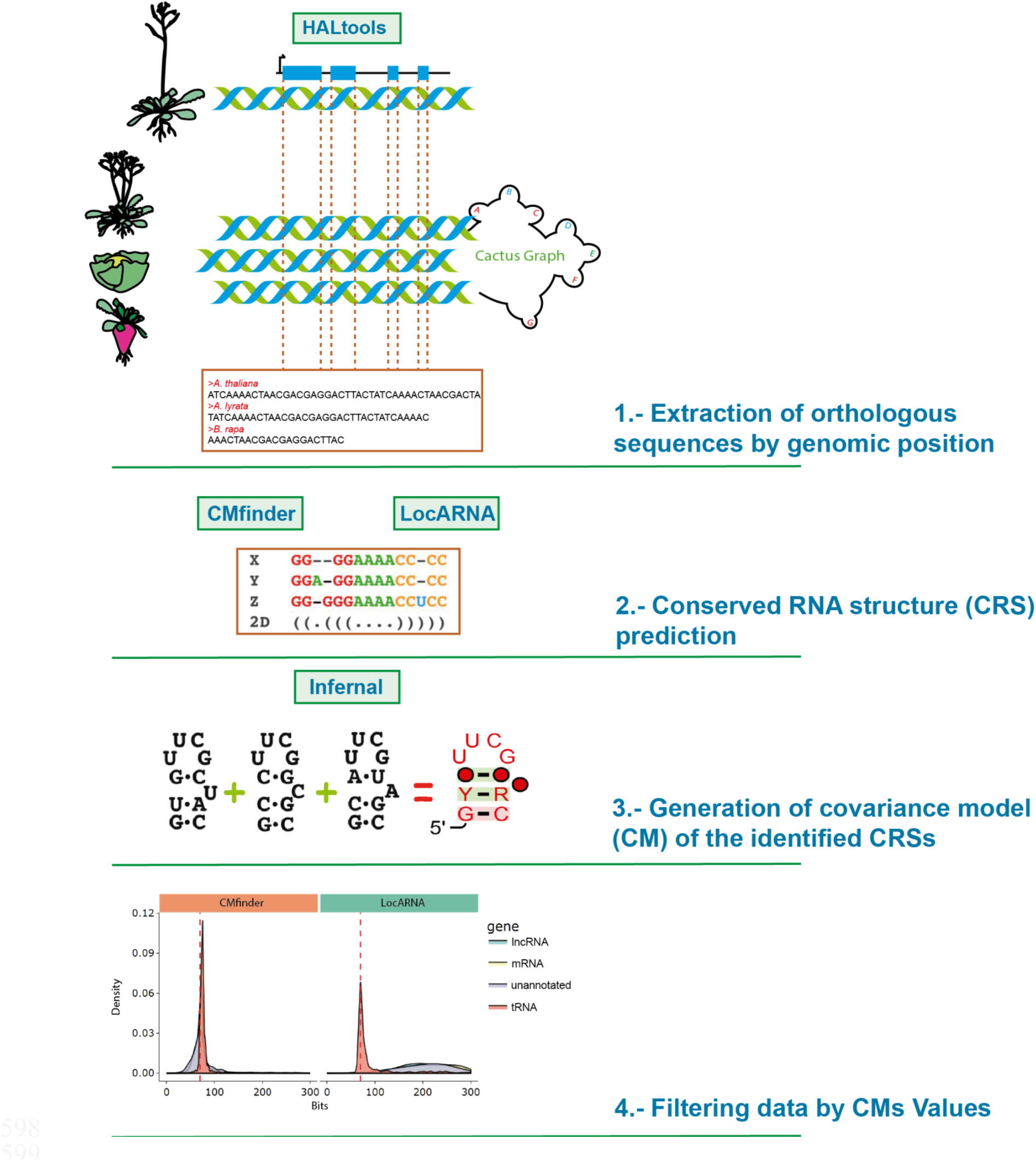
– Methodology for Identifying Conserved RNA Motifs. Diagram illustrating the process used to identify conserved RNA motifs. The first step involves extracting the exons of each gene from a genomic alignment and storing them in FASTA format (1). In the next step, conserved structural motifs are searched for and stored in Stockholm format (2). Next, covariance models are generated (3), providing us with conservation values for the identified motifs. Finally, values are filtered to identify a final set of CRSs (4).

### CRSs prediction

To predict CRSs, we realigned the sequences extracted in the previous step using two structure-aware programs: CMFinder and LocARNA. This realignment step is important as the original alignment of the regions is only based on sequence homology, which could lead to errors in the prediction of the conservation of secondary structures. The prediction of conserved structural motifs and a sequence realignment were done simultaneously (Figure 1).

The first method, CMfinder, is a tool used for the prediction of RNA structural motifs. It uses a heuristic search for structural motifs, combining folding energy values and covariance models, which can be used at the genome scale (Yao et al. 2006). CMfinder realigns the input alignment blocks, using a statistical method to favor the alignment of the possible structure, improving the accuracy of the prediction but at a high computational cost (Smith et al. 2013). Due to its high computing time, it is limited to finding small conserved RNA framework motifs of around 200 bases (Seemann et al. 2017; Torarinsson et al. 2008). CMfinder reports ∼79% of accuracy in its prediction of structural motifs compared to structures already established in Rfam (Yao et al. 2006).

The second method, LocARNA, is an alignment algorithm based on sequence similarity and secondary structure; it works by simultaneously calculating the probability of coincidence of sequence and structure (Will et al. 2007). It has been noted that the accuracy of the structural prediction increases when aligned simultaneously to the sequence, increasing the quality of the alignment in sequences that have little identity compared to alignments based on sequence alone (Hofacker and Lorenz 2014). We used locARNA-p, which is an improvement on locaRNA in terms of speed and prediction precision (Will et al. 2012). locARNA-p reports an accuracy of ∼87% using a specialized database to measure the efficiency of structural aligners (Will et al. 2012; Wilm et al. 2006). Unlike CMfinder, locARNA-p can process larger structural motifs of around 1,000 bases.

### Generation of a covariance model (CM) of the identified CRSs

In the third phase of the analysis, we created probabilistic models called covariance models (CMs) to describe the relationships between nucleotide positions in RNA sequences and the structure of RNA molecules. These models enable the inclusion of compensatory substitutions in the nucleotides that maintain the conserved structure in the structural prediction. The statistical combination of sequence and structure homology in the CMs enables the classification and comparison of RNA structural motifs (Nawrocki et al. 2009, 2015; Nawrocki 2014). CMs are used by the Rfam database, which is a reference database used to annotate functional RNA motifs. The database contains a vast collection of CMs from various RNA families (Nawrocki et al. 2015). To generate the CMs, we use the Infernal program v1.1.2 (Nawrocki et al. 2009), which is based on stochastics with context-free grammars (SCFGs) to create CMs of the alignments with structural prediction.

To compare both conservation methods and compare the descriptive values of our identified structures, we use the results of Infernal (Nawrocki et al. 2009), from which we extracted a series of descriptive values of the structure including number of species included in the CRSs motif, consensus positions on alignment, size of alignment, number of paired bases and entropy values for the CM. Of these descriptive values, the most important is the value in CM bit-score, which tells us the amount of information given to us by sequence conservation plus structure by alignment position; this is followed by the value of bits that would be generated by only the sequence with a hidden Markov Model (HMM) without considering the structure. With the difference in these values, we can know the information gained by the structure in the alignment (Figure S1-F).

### Data filtering

To find the conserved structures, we followed a set of filters. Initially, we only considered structures that were present in two or more species in the genomic alignment. Then, we filtered the structures based on their average information level, keeping only those with a level above 66.68 bits. This value was obtained from the information level of tRNAs (as shown in Figure 2), and we used it to ensure that we only included CRSs with a level of information that was at least as high as that of tRNA structures. This filtering step also helps to remove any insignificant CRSs.

**Figure 2.**
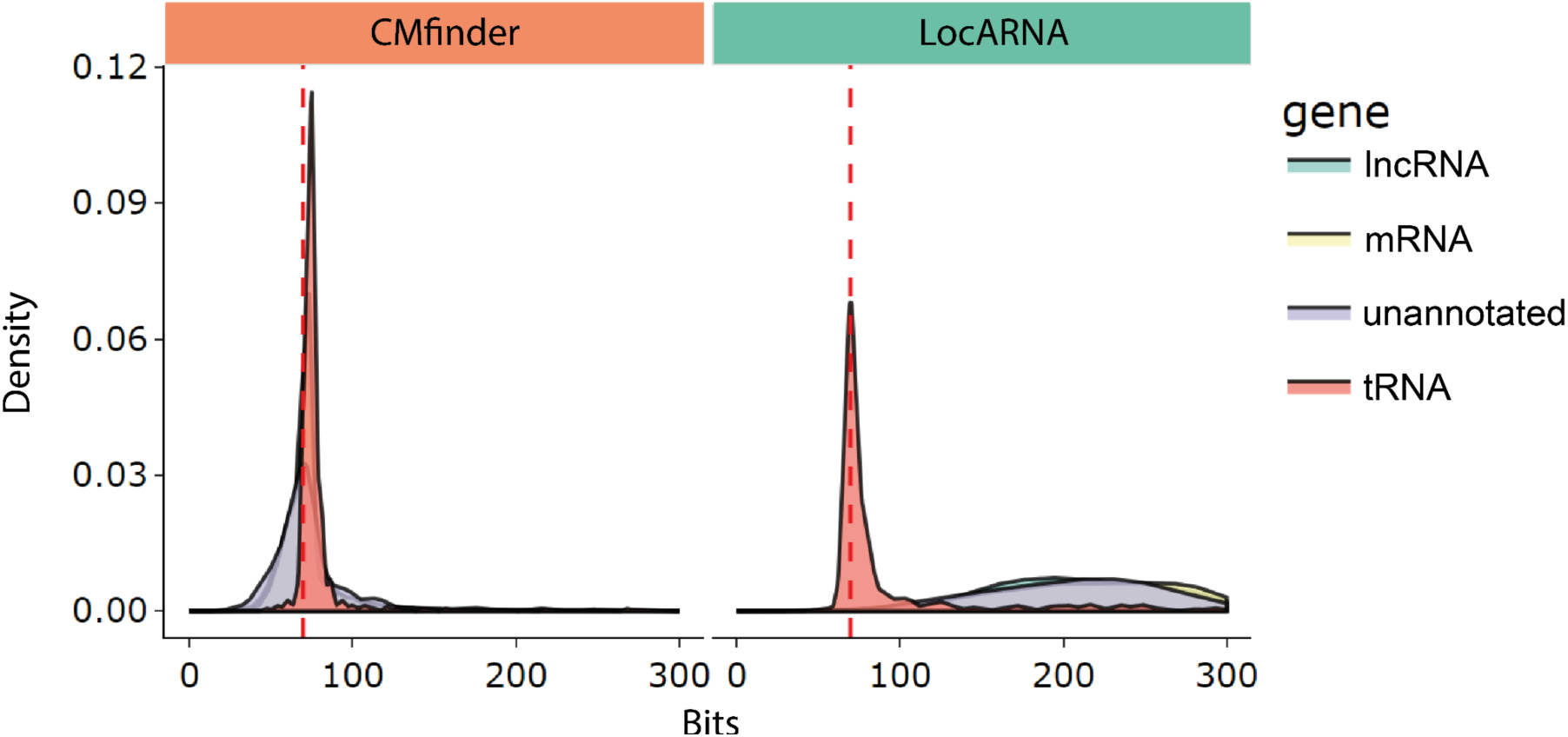
– Density graph of the amount of information in bits of the covariance model for each method. In each of the graphs the 4 gene biotypes used in the analysis and their amount of information are shown. The red dotted line represents the average mean bit value of the tRNAs (66.68 bits) which is the cut-off value used to decide if a structure has a significant amount of information.

**Figure 3.**
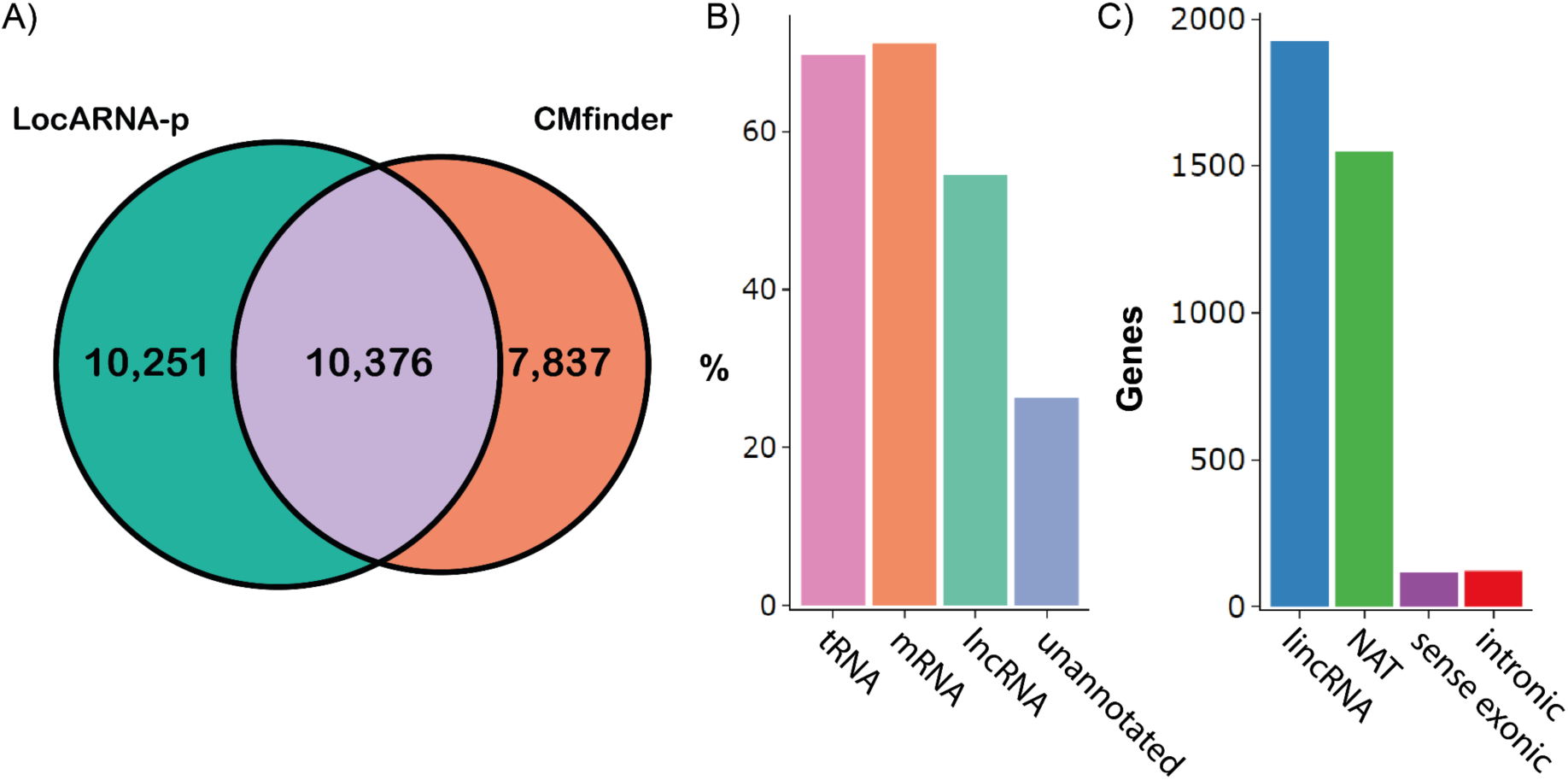
– Predicted CRSs. A) Venn diagram of the predicted CRSs in each method. B) Percentage of genes that have CRSs and unannotated regions of the genome compared to lncRNAs. C) lncRNAs containing CRSs divided into the different biotypes of lncRNAs.

**Figure 4.**
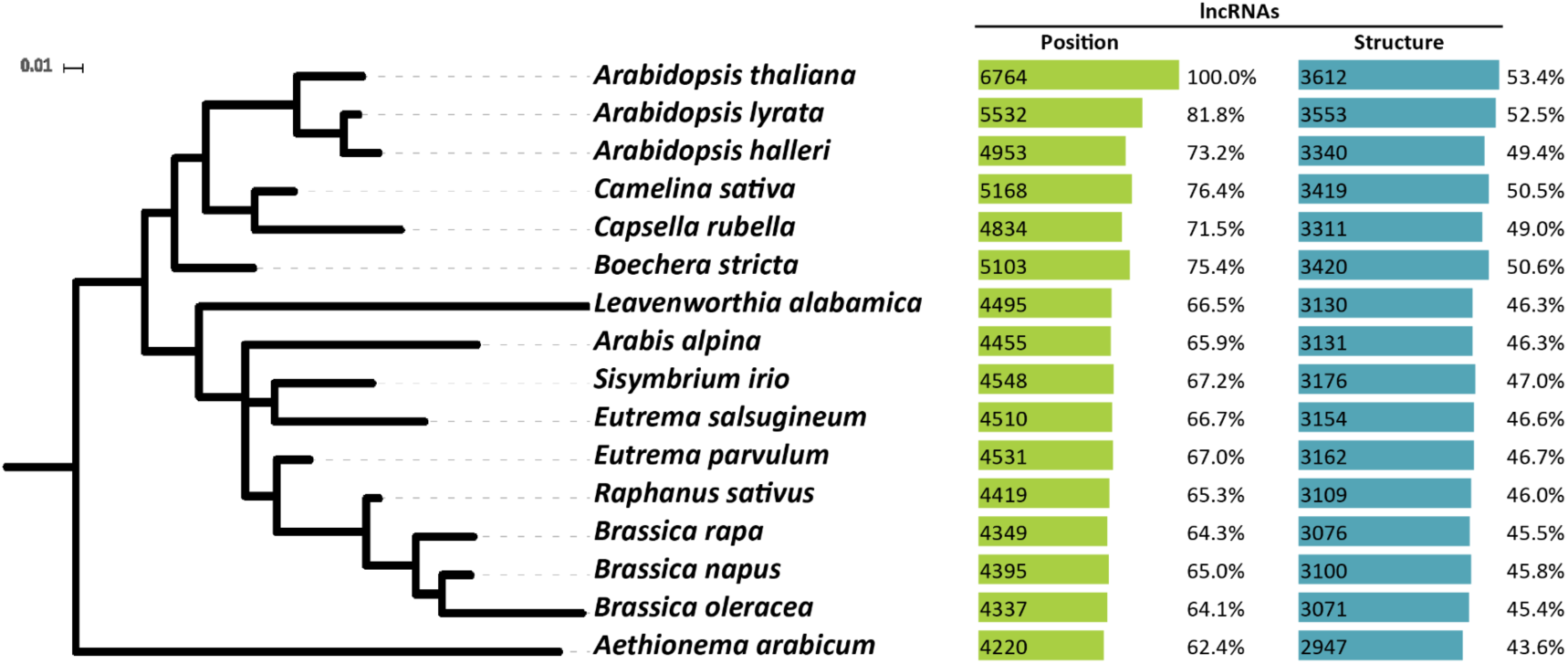
– Conservation by position and structure of the lncRNAs in each of the 16 species of *Brassicaceae*. The number and percentage of lncRNAs identified as conserved by position (green, left) and structure (blue, right) are shown. Phylogenetic tree scale is in changes per site.

**Figure 5.**
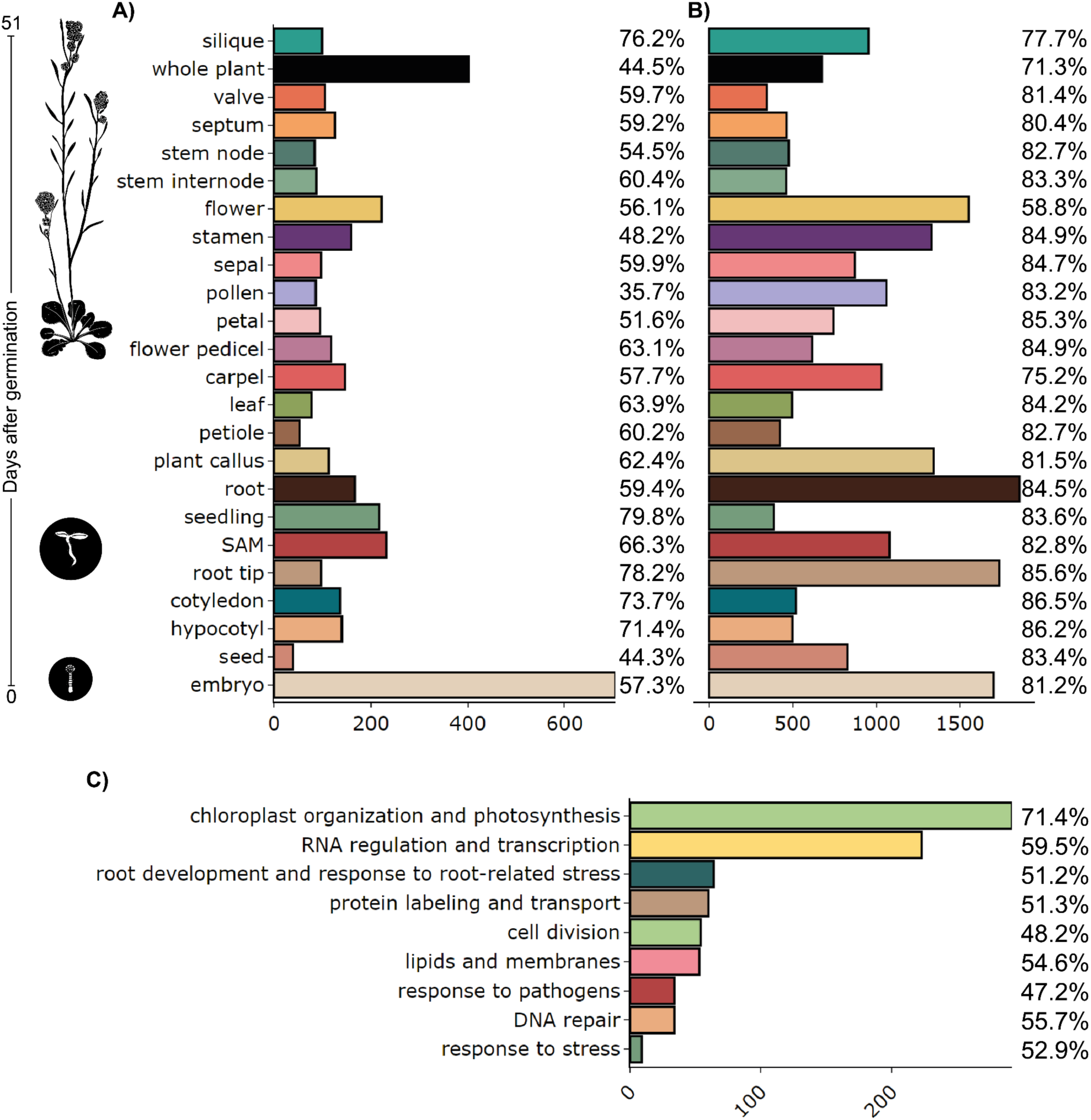
– Distribution of CRSs in tissue-specific lncRNAs and in lncRNAs associated with functional categories. A) Number of lncRNAs that have CRSs and are tissue and developmental stage-specific B) mRNAs with CRSs and are tissue-specific. C) lncRNAs identified by co-expression in functional categories.

**Figure 6.**
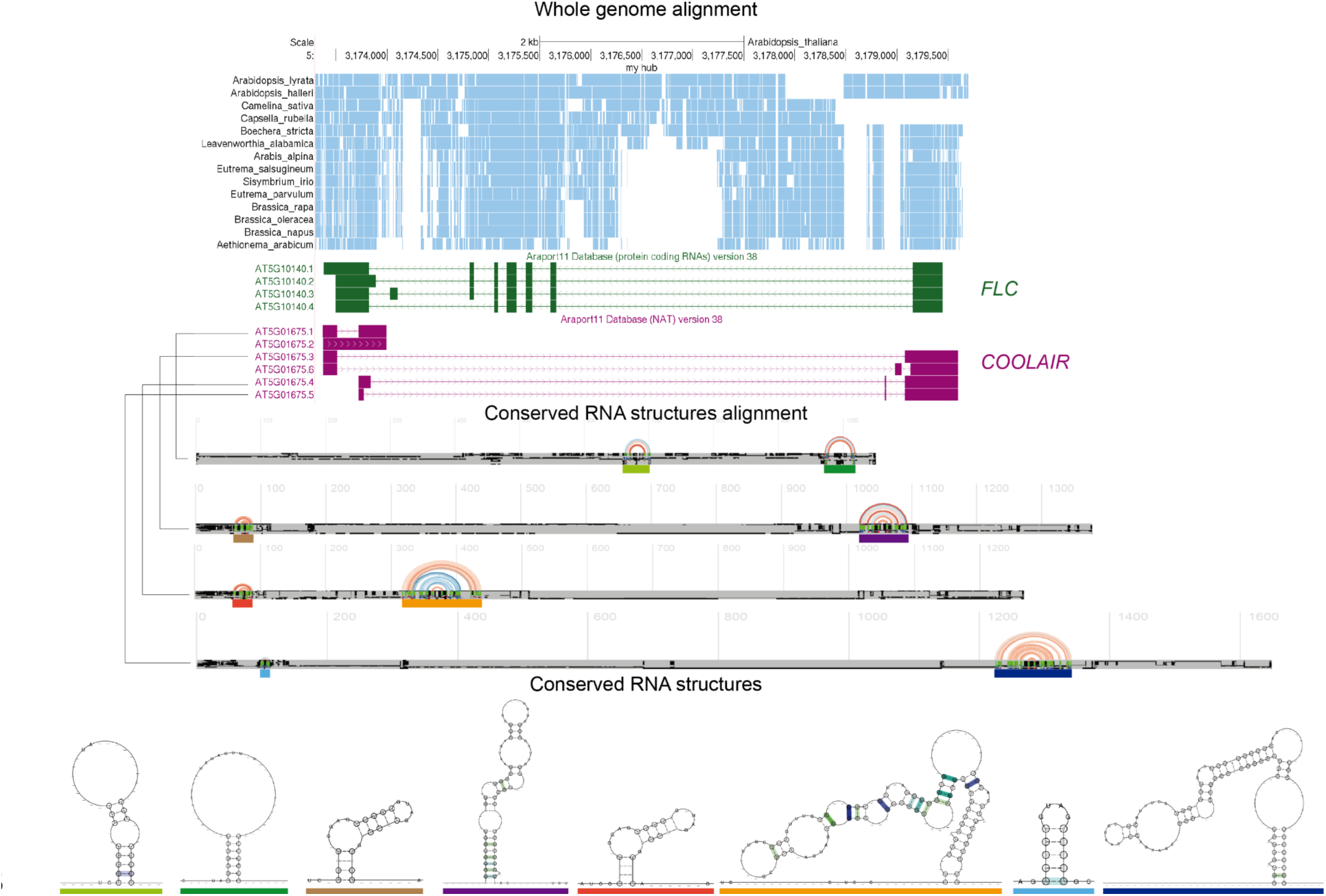
– Structural conservation prediction in *COOLAIR*. The window of genome alignment is displayed in the UCSC genome browser. Exons and introns of the annotated genes in the window. CRS alignment drawing with arcplot R-chie (The color of the arcs represents the covariance levels of the paired bases, red higher covariance values blue lower covariance values). Conserved secondary structures using RNAplot.

To assess how many of the predicted structures have a covariance above that expected phylogenetically, we assessed the significance for each position with the program R-scape v2.0.0.j (Rivas et al. 2017) on the Stockholm format alignments. To consider a positive result in this test, We only retained structures with more than five paired bases with covariance where the observed covariance was greater than phylogenetically expected according to R-scape.

### Visualization and handling of CRS coordinate

To compare the results of structural conservation with the previous types of conservation (position conservation and splicing conservation), we identified the genomic coordinates of CRSs and represented them graphically in a genome viewer track. To obtain the genomic coordinates we used Blat (-q=rna –t=dna –out=psl) (Kent 2002) using the sequence of the CRSs as a query to determine its position within the genome. This allowed us to represent CRSs in their spliced form. The UCSC Genome Browser can be used to visualize the different types of conservation in one gene, and ability to juxtapose annotations of different types of conservation for comparison (Kuhn et al. 2013).

## Acknowledgements

This work was funded in part by Consejo Nacional de Ciencia y Tecnologia (CONACYT PhD Scholarship 338379 [J.A.C.-G.], and by a Royal Society Newton Advanced Fellowship (NAF\R1\180303) awarded to S.L.F.-V.

