## Supplemental Material for "Evolutionary conservation of secondary structures in the lncRNAs of plants"

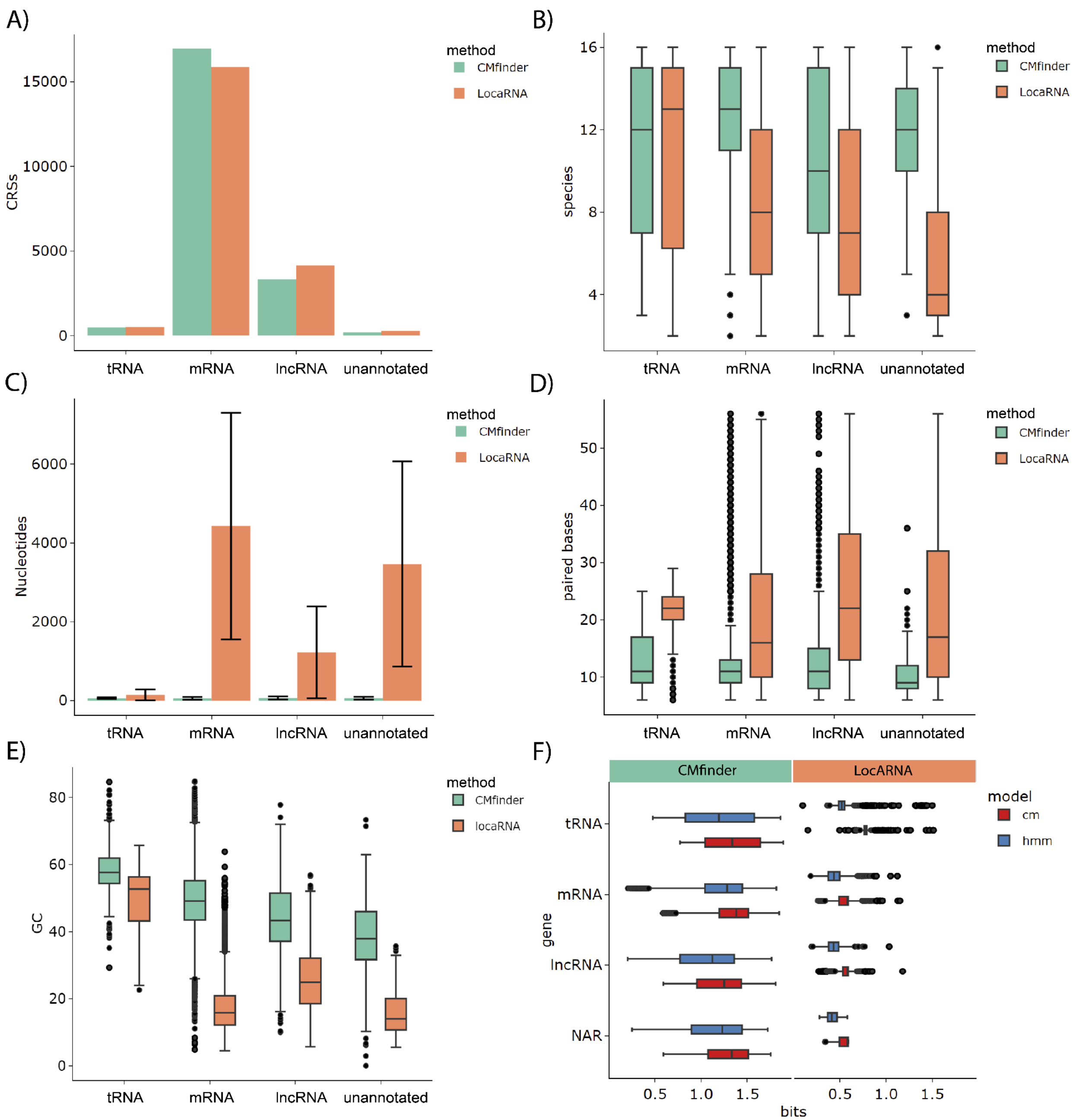


**Figure S1 - Descriptive values of CRSs by the method** A) Number of CRRs identified in each of the biotypes, B) Number of species included in the CRSs motif, C) Size of the CRSs alignment including gaps, D) Number of paired bases in the CRSs, E) GC percentage of the CRSs and F) Information gain of the covariance model over the hidden Markov model.

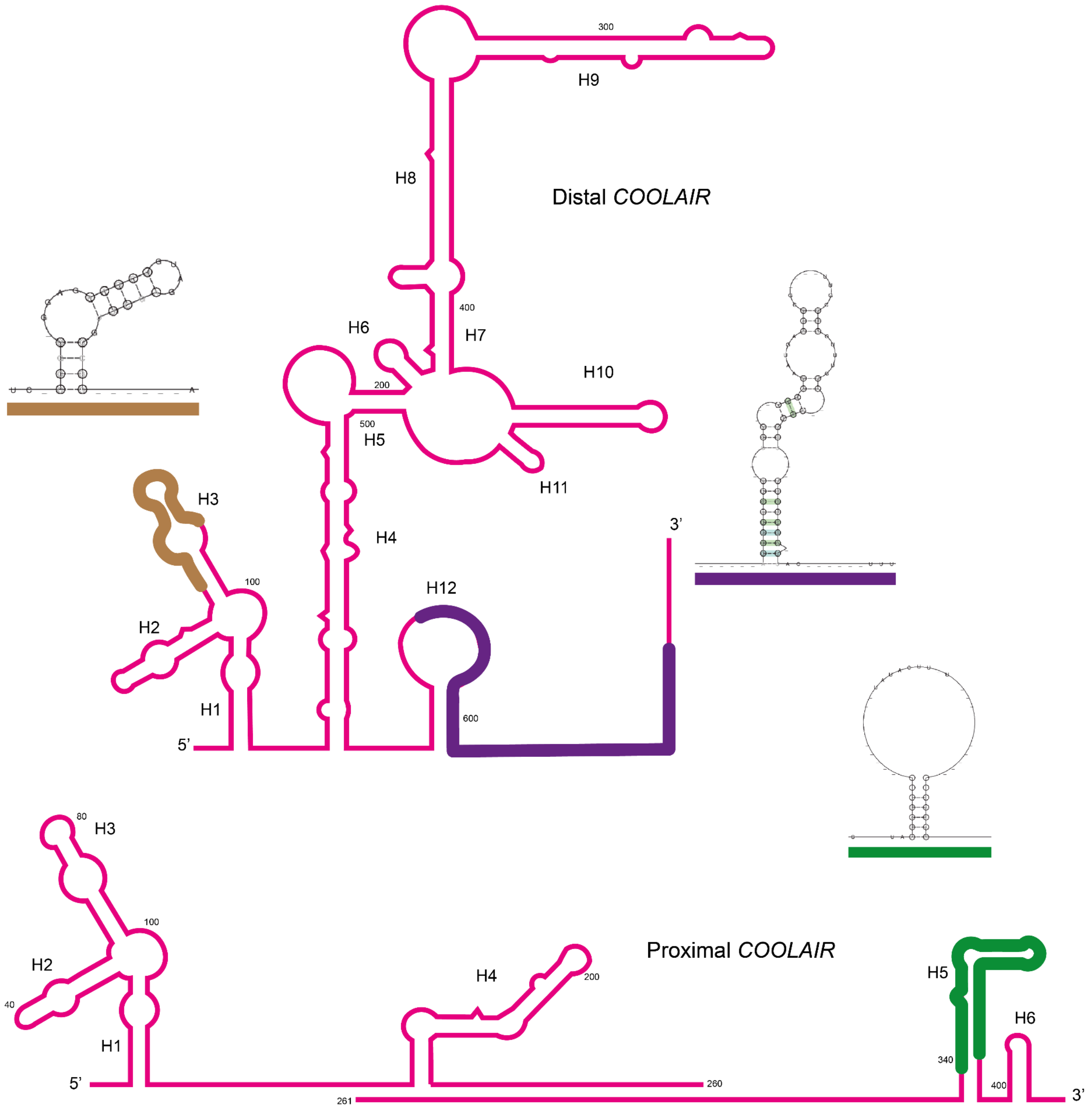


**Figure S2 - Comparison of CRSs with *COOLAIR* structure prediction, figure modified from Hawkes et al 2016.** The predicted structure of each CRS is shown in the same color (brown, purple or green) outside the previously published structures and highlighted in the same color within previous predictions both for distal *COOLAIR* (top) and proximal *COOLAIR* (bottom).
